# Spatiotemporal control of subcellular O-GlcNAc signaling using Opto-OGT

**DOI:** 10.1101/2024.05.12.593740

**Authors:** Qunxiang Ong, Rachel Lim, Cameron Goh, Yilie Liao, Sher En Chan, Crystal Lim, Valerie Kam, Jerome Yap, Tiffany Tseng, Reina Desrouleaux, Loo Chien Wang, Siok Ghee Ler, Siew Lan Lim, Sunyee Kim, Radoslaw M Sobota, Anton M. Bennett, Weiping Han, Xiaoyong Yang

## Abstract

The posttranslational modification of intracellular proteins through O-linked β-N-acetylglucosamine (O-GlcNAc) is a conserved regulatory mechanism in multicellular organisms. Catalyzed by O-GlcNAc transferase (OGT), this dynamic modification plays an essential role in signal transduction, gene expression, organelle function, and systemic physiology. Here we present Opto-OGT, an optogenetic probe that allows for precise spatiotemporal control of OGT activity through light stimulation. By fusing a photosensitive cryptochrome protein to OGT, Opto-OGT can be robustly and reversibly activated with high temporal resolution by blue light and exhibits minimal background activity without illumination. Transient activation of Opto-OGT results in mTORC activation and AMPK suppression which recapitulate nutrient-sensing signaling. Furthermore, Opto-OGT can be customized to be localized at specific subcellular sites. By targeting OGT to the plasma membrane, we demonstrate downregulation of site-specific AKT phosphorylation and signaling outputs in response to insulin stimulation. Thus, Opto-OGT is a powerful tool to define the role of O-GlcNAcylation in cell signaling and physiology.

---

Intracellular signaling processes utilize reversible posttranslational modifications (PTMs) to modulate the fate and function of individual proteins. Among the growing list of PTMs, O-GlcNAcylation represents an important intracellular glycosylation event. It involves the attachment of O-linked β-N-acetylglucosamine (O-GlcNAc) moieties onto serine or threonine residues of proteins in various cellular compartments, including the nucleus, mitochondria, plasma membrane, and cytoplasmic space.^1–3^. *O*-GlcNAcylation is controlled by a single pair of enzymes, O-GlcNAc Transferase (OGT) and O-GlcNAcase (OGA), which drive the addition and removal of O-GlcNAc moieties, respectively^4–8^. Functionally, O-GlcNAcylation is primarily thought to regulate diverse cellular processes in response to nutrient availability or various stress conditions^9,10^. Dysregulation of O-GlcNAc cycling has been implicated in the pathogenesis of various diseases, such as cancer, diabetes and neurodegeneration^11–15^. Hence, therapeutic strategies aimed at modulating O-GlcNAc signaling have been explored for degenerative diseases^16^. For instance, clinical trials are underway for OGA inhibitors such as MK-8719^17^ and ASN120290^18^ for the treatment of progressive supranuclear palsy and Alzheimer’s disease, respectively.

The combinatorial modification of proteins with multiple PTMs generates a dynamic ‘code’ that controls protein function^9,19,20^. However, the development of therapeutic strategies to modulate cellular O-GlcNAc levels has been challenging due to a limited understanding of O-GlcNAc signaling dynamics as part of the PTM code^3^. OGT and OGA are dynamically regulated in cells over space and time, leading to different outcomes depending on their interacting partners, subcellular localization and duration of interaction^10,21^. The complexity of studying the role of O-GlcNAcylation is further compounded by the intricate network of signaling events that arise when several signaling pathways can be simultaneously activated in physiological settings. For instance, mTORC, AMPK and OGT signaling modules are involved in glucose sensing and nutrient deprivation. Thus, precise manipulation of OGT activity is critical for dissecting the cellular processes regulated by O-GlcNAc modification, thereby improving our understanding of the PTM code in regulating human health and disease.

Intracellular optogenetics has been shown to be a powerful platform for dissecting complex signaling modalities with high spatiotemporal precision, as these tools are reversible, possess little background activity, and respond rapidly to light stimulus^22–24^. Genetically-encoded light responsive proteins, such as light, oxygen or voltage domains (LOV)^25^, cryptochrome^26^ (CRY2) and phytochrome^27^ have been utilized to control protein location, activity and association. In particular, optogenetic tools to modulate the activity of kinases such as tyrosine kinases^28,29^, mitogen-activated protein kinase (MAPK)^30,31^ and Akt^32^ have been developed using protein oligomerization and localization strategies. While optogenetic tools have mainly been utilized to probe kinase signaling, transcriptional activities^33–35^ and GTPases^36–38^, we hypothesize that the approach can also be used to control OGT activation and allow us to probe for signaling transmission by O-GlcNAcylation.

Here, we report the generation of light-activated OGT driven by the oligomerization and predicted conformational change of cryptochrome-tagged OGT, which we refer to as ‘Opto-OGT.’ Upon light activation, Opto-OGT increases O-GlcNAc levels in cells globally and interacts with similar protein partners as OGT. Opto-OGT is tunable and reversible, allowing for precise control of OGT activation kinetics and signaling dynamics. We demonstrate the utility of Opto-OGT in studying glucose sensing in HEK293T cells, showing that a 15-minute activation of OGT results in similar changes to cellular signaling as glucose replenishment. Additionally, we leverage the hetero-dimerization properties of CRY2 and CIBN (N-terminus truncation of CIB1, a CRY2 interaction partner) to recruit OGT to specific subcellular compartments, such as the mitochondria and plasma membrane. We demonstrate that OGT recruitment to the plasma membrane downregulates AKT signaling. Overall, Opto-OGT represents a novel tool for manipulating the spatiotemporal aspects of O-GlcNAc signaling, providing insights into the role of O-GlcNAcylation in cellular physiology and disease.

## Results

### Design principles of a light-activated OGT toolkit, Opto-OGT

To achieve precise spatiotemporal control of OGT activity, we propose the installation of a blocker at the UDP-GlcNAc substrate entry site of OGT, leading to an inactivated form. This inactivated form can then undergo controlled conformational changes in response to a stimulus, resulting in the release of the substrate entry site and subsequent activation of OGT (**Figure 1A**). We reasoned that an optogenetic system driven by CRY2 conformational change, here referred to as the Opto-OGT system, could control OGT enzymatic activity in a light-dependent fashion. The full-length human OGT is fused to the fluorescent reporter mCherry at the C-terminus and is subsequently tagged with the crytochrome construct. In this design, OGT-mCherry-CRY2 is expected to be localized in the cytosol as discrete proteins in the dark, with the CRY2 acting as the blocker towards OGT substrate entry site. Upon blue light excitation, the CRY2 will undergo conformational changes and release the OGT substrate entry site thus activating it (**Figure 1A**). The design was supported by AlphaFold simulation^39^, which predicted the crytochrome resides at the substrate binding pocket of OGT in dark state (**Figures 1B and S1**). We also examined the heterodimerization of CRY2 and OGT as two separate proteins *in silico* with both ClusPro^40^ (**Figure S2A**) and AlphaFold (**Figure S2B-E**) for unbiased analysis. Similarly, CRY2 in the dark state is expected to cluster at the substrate entry site of OGT without any tethering effects.

**Figure 1:**
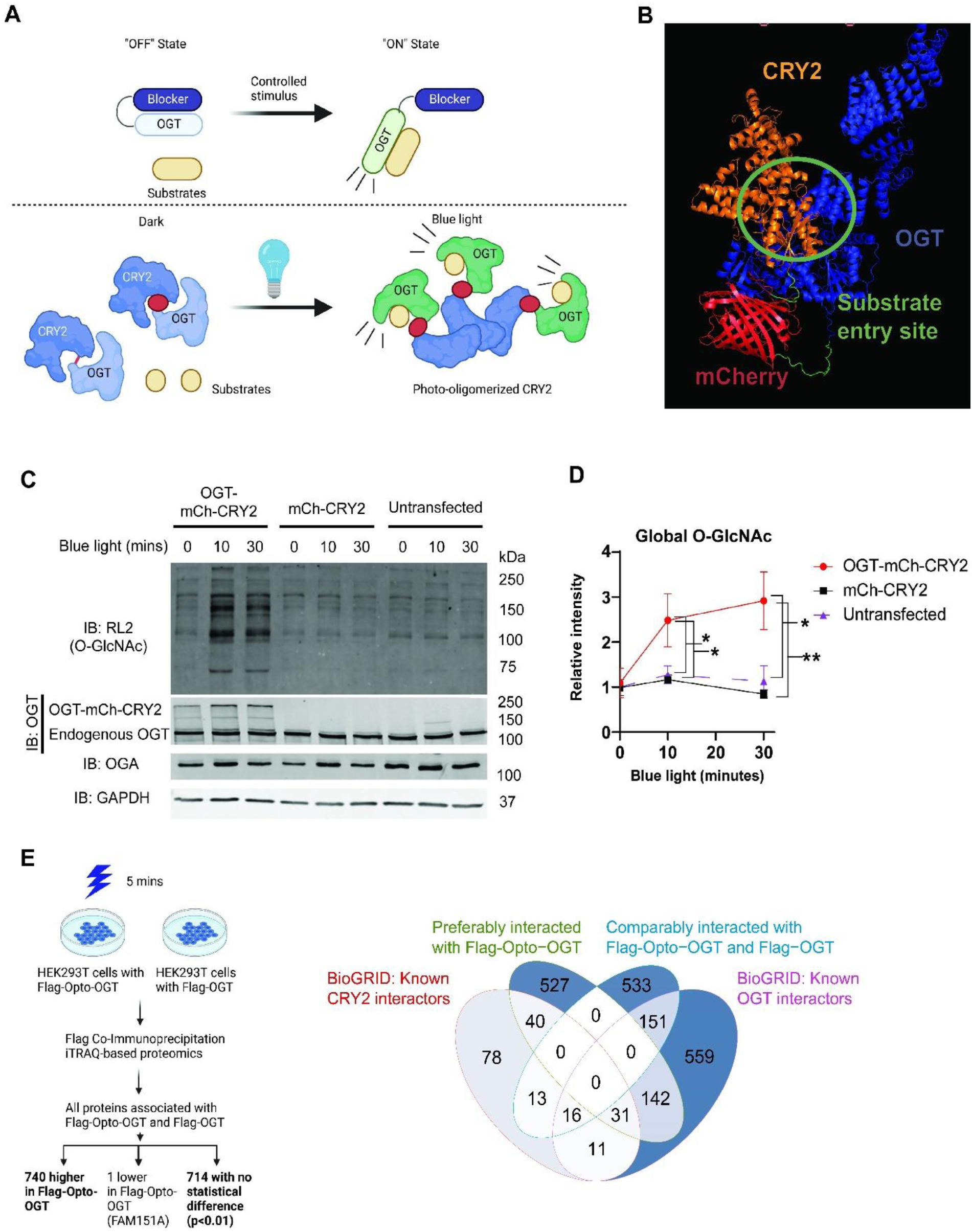
Design and characterization of Opto-OGT. (A) The ideal OGT signaling toolkit is illustrated in the top panel, where OFF-state OGT does not have access to substrates while ON-state OGT, under a controlled stimulus, gains access to substrates. Schematic of light-controlled OGT oligomerization and activation is shown in the bottom panel. OGT-mCherry-CRY2 is expressed as a fusion protein and CRY2 blocks access of substrates to OGT in dark. Upon blue light exposure, CRY2 oligomerization results in conformation change of OGT-mCherry-CRY2 and local proximity aggregation of OGT leading to its activation. (B) AlphaFold simulation of OGT-mCherry-CRY2 in the dark state reveals that CRY2 blocks access to OGT substrate binding pocket. (C) Representative Western blots of RL2 (Global O-GlcNAc levels), OGT, OGA and GAPDH of HEK293T cells transfected with OGT-mCherry-CRY2 or mCherry-CRY2, or untransfected ones with or without blue light. (D) Quantification (n=3) of normalized global O-GlcNAc levels relative to GAPDH for Western blots depicted in (B). (E) Mass spectroscopy of proteins associated with Flag-Opto-OGT or Flag-OGT demonstrates that Flag-Opto-OGT retains protein interactions with similar proteins as Flag-OGT. Venn diagram reveals the number of proteins that similarly or preferably interacted with Flag-Opto-OGT and/or Flag-OGT. Data are presented as mean ± s.d. Statistical analysis employed is Student’s t test with * indicating p<0.05 and ** indicating p<0.01.

### Light-driven activation of Opto-OGT results in local increases in O-GlcNAc levels

We first asked if light-driven activation of Opto-OGT results in an increase in overall O-GlcNAc levels in the cells. HEK293T cells were transfected with either OGT-mCherry-CRY2 or mCherry-CRY2, or did not undergo transfection. These cells were subjected to blue light illumination for 0, 10 or 30 minutes using a home-made LED array. We found that only OGT-mCherry-CRY2 increased overall O-GlcNAc levels upon blue light illumination, whereas the O-GlcNAc levels were comparable between OGT-mCherry-CRY2 or mCherry-CRY2-transfected and untransfected cells at 0 minute (**Figures 1C and 1D**). Expression of OGT-mCherry-CRY2 did not result in a significant increase in total OGT and OGA protein levels compared to the controls (**Figure S3A**). These results are corroborated by the immunofluorescence staining of COS7 cells showing the presence of O-GlcNAc puncta (indicated by RL2 antibody) in the areas of OGT-mCherry-CRY2 oligomerization under blue light (**Figure S4**). In addition, the light supplied at 200 µW/cm^2^ did not activate stress-responsive MAPK activity, confirming that blue light illumination has no detrimental effect on the cells (**Figure S5A**).

### OGT interactome is captured similarly by Opto-OGT

To confirm that Opto-OGT functions similarly as endogenous OGT in the activated state, we asked if the identity of proteins that interact with Opto-OGT is similar to that with OGT. In this regard, we subjected HEK293T cells transfected with FLAG-tagged Opto-OGT to five minutes of blue light and performed co-immunoprecipitation experiments to capture the FLAG-tagged proteins. The FLAG-tagged Opto-OGT was confirmed for its ability to increase O-GlcNAc levels upon blue light stimulation. (**Figure S3B**). Comparing the lists of proteins associated with FLAG-Opto-OGT and FLAG-OGT, we found that 740 proteins are more significantly associated with Flag-Opto-OGT while 714 proteins are similarly associated to both Flag-Opto-OGT and Flag-OGT. (**Figure 1E**). We next compared the lists of proteins with previously known proteins that bind to CRY2 or OGT according to the BioGRID database. We found proteins that are preferably bound to Flag-Opto-OGT share a limited subset associated with CRY2 (40) while the majority of the proteins are known OGT interactors (142+31). Similarly, the list of proteins that are similarly associated with both Flag-Opto-OGT and Flag-OGT contains a significant number of known OGT interactors (151+16) (**Figure 1E and Table S3**). This mass spectrometry analysis thus reveals the nature of the Opto-OGT interactome to be similar to that of OGT.

### Opto-OGT activity is reversible and tunable

After confirming the functionality of Opto-OGT in driving O-GlcNAcylation, we further characterized the reversibility and tunability of this tool. We first examined the temporal kinetics of the oligomerization of the fusion protein. The results showed that OGT-mCh-CRY2 formed aggregates within 5 minutes under blue light (**Figures 2A and 2B, Movie S1**). Following 5 minutes of blue light exposure, the cells were subjected to darkness for 20 minutes. We found that the number of oligomers decreased to the basal levels within 5 to 7 minutes (**Figure 2B**). Reactivation of OGT-mCh-CRY2 by blue light led to the formation of oligomers to the same extent as the previous cycle (**Figure 2B**). We next tested if the changes in Opto-OGT oligomerization translate to respective changes in O-GlcNAc levels of the cell. Indeed, 5 minutes of blue light increased O-GlcNAc levels, while it took about 2 hours to reset the O-GlcNAcylation levels to the baseline. Reactivation of Opto-OGT oligomerization for 10 minutes was also sufficient to raise the overall O-GlcNAc level of the cell, indicating the reversibility of the system (**Figure 2C**).

**Figure 2:**
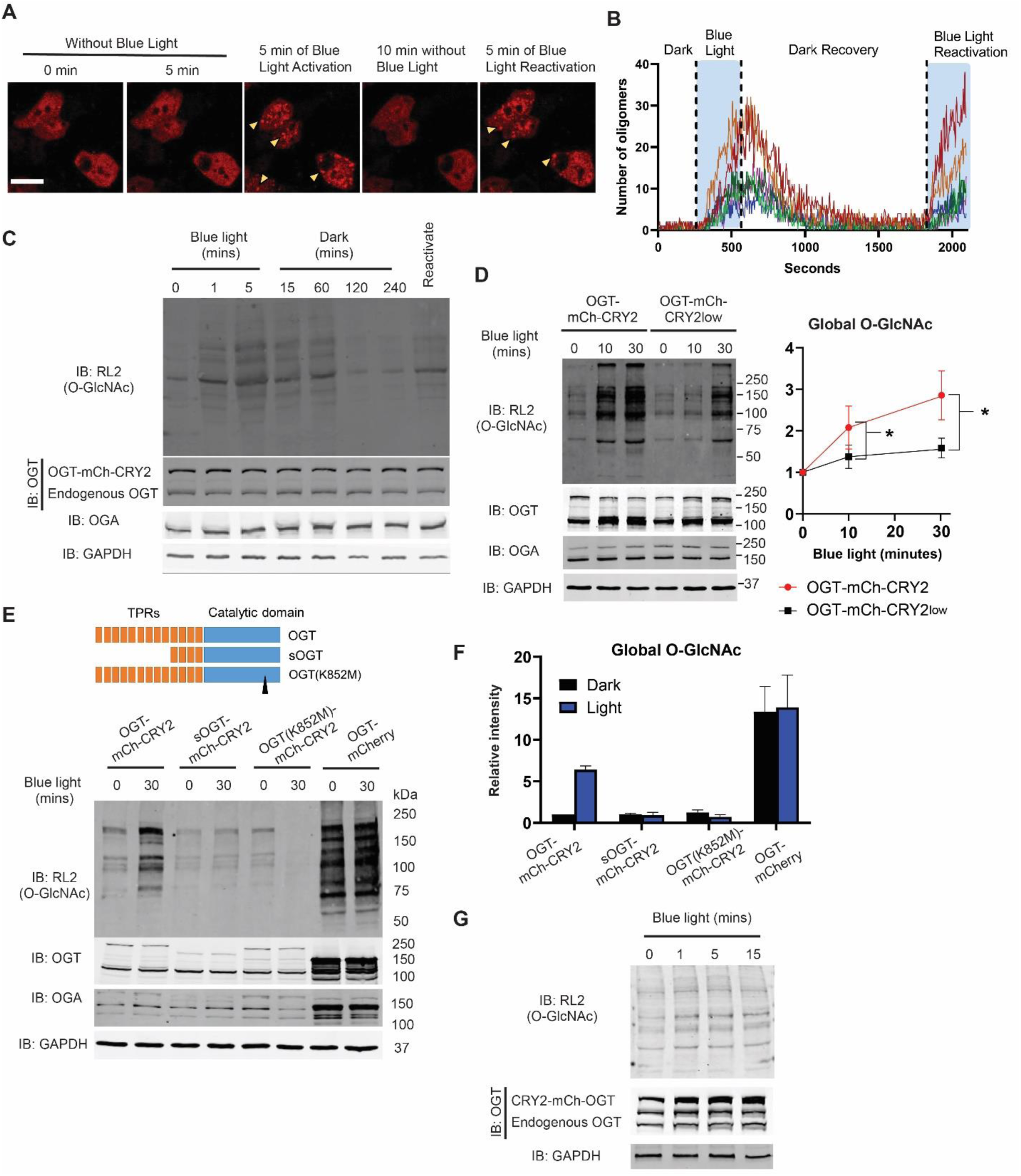
Opto-OGT is tunable and reversible, and further characterization of Opto-OGT reveals the necessary requirements for it to work. (A) Representative images of HEK293T cells transfected with OGT-mCherry-CRY2 and undergoing dark-blue light cycles. The yellow arrows indicate the observation of puncta, indicating the formation of oligomers. Scale bar = 20 µm (B) Quantification of oligomers in cells (n=21) undergoing dark-blue light cycle (C) Representative Western blots of RL2 (Global O-GlcNAc levels), OGT, OGA and GAPDH of HEK293T cells transfected with OGT-mCherry-CRY2 and exposed to a series of conditions as labelled. (D) Representative Western blots of RL2 (Global O-GlcNAc levels), OGT, OGA and GAPDH of HEK293T cells transfected with OGT-mCherry-CRY2 or OGT-mCherry-CRY2low, with or without blue light. Quantification (n=3) of normalized global O-GlcNAc levels relative to GAPDH for Western blots depicted. (E) Schematic of the different OGT proteins studied in the Western blot. Tetratricopeptide repeats and catalytic domain of the different proteins are represented. Representative Western blots of RL2 (Global O-GlcNAc levels), OGT, OGA and GAPDH of HEK293T cells transfected with OGT-mCherry-CRY2 or mCherry-CRY2, or untransfected ones with or without 30 minutes of blue light. (F) Quantification (n=3) of normalized global O-GlcNAc levels relative to GAPDH for Western blots depicted in (E). (G) Representative Western blots of RL2 (Global O-GlcNAc levels), OGT, OGA and GAPDH of HEK293T cells transfected with CRY2-mCherry-OGT with or without blue light. Data are presented as mean ± s.d. Statistical analysis employed is Student’s t test with * indicating p<0.05 and ** indicating p<0.01.

To test whether the increase in global O-GlcNAc levels could be tuned by different rates of CRY2-induced conformational changes, we generated OGT-mCh-CRY2^low^ which has a lower rate of oligomerization^41^. We argue that OGT-mCh-CRY2^low^ changes its conformation more slowly and releases active OGT at a slower rate. Under blue light, the rate of increase in O-GlcNAcylation was higher in the cells expressing OGT-mCh-CRY2 compared to those expressing OGT-mCh-CRY2^low^ (**Figure 2D**). These results demonstrate that the Opto-OGT system is a reversible, tunable, and robust optogenetic tool to control O-GlcNAc signaling.

We next determined which regions in Opto-OGT are critical for light-controlled OGT activity by introducing a truncation or point mutation into the Opto-OGT sequence. After 30 min of blue light exposure, the full-length OGT fused with mCherry-CRY2 markedly elevated cellular O-GlcNAcylation (**Figure 2E**). In contrast, the short OGT (sOGT) and catalytically dead OGT (K852M) failed to elevate global O-GlcNAc levels (**Figures 2E and 2F**). These results indicate that both the tetratricopeptide repeat (TPR) domain and the catalytic domain are required for Opto-OGT function. On the other hand, overexpression of OGT-mCherry without the CRY2 domain resulted in constitutively higher O-GlcNAc levels despite a compensatory increase in OGA expression (**Figures 2E and 2F**). Titrated levels of OGT-mCherry overexpression are proportional to the subsequent global O-GlcNAc increase (**Figure S5B**). These results indicate that light-driven activation of the full-length opto-OGT increases cellular O-GlcNAcylation levels. We also asked if the order of the fusion protein is critical. As such, we constructed CRY2-mCherry-OGT and activation via blue light causes a modest increase in the O-GlcNAcylation level of the cell, indicating that the positioning of the cryptochrome with respect to OGT may affect the effectiveness of the optogenetic construct (**Figure 2G**).

### Analysis of AMPK, mTORC and O-GlcNAc signaling pathways upon glucose level changes

Glucose concentration is continually sensed by cells to allow adequate responses to any changes in its availability^42^. Such changes have been proposed to be sensed by AMPK, mTORC and OGT, and could lead to cellular responses such as fatty acid synthesis, autophagy and proliferation^43,44^ (**Figure 3A**). We first subjected cells to 3 hours of treatment at 0 mM glucose, and then subject them to 25 mM glucose for 15, 30 and 60 minutes, respectively. Global O-GlcNAc levels increased within 15 minutes of glucose concentration change, while phospho-acetyl coenzyme A (ACC) decreased within the same timeframe (**Figures 3B and 3E**). Notably, the upstream kinase of ACC, AMPK, did not post any significant changes in its phosphorylation level (**Figures 3B and 3E**). Phosphorylation of mTORC signaling components including mTOR and p70S6K did not show significant increases in their phosphorylation levels (**Figure S6A**), whereas phosphorylation of S6 increased at 30 and 60 minutes (**Figures 3B and 3E**). Notably, O-GlcNAcylation of AMPK showed a prominent increase at 15 and 30 minutes, which could potentially modulate AMPK function independent of its phosphorylation (**Figure 3B**).

**Figure 3:**
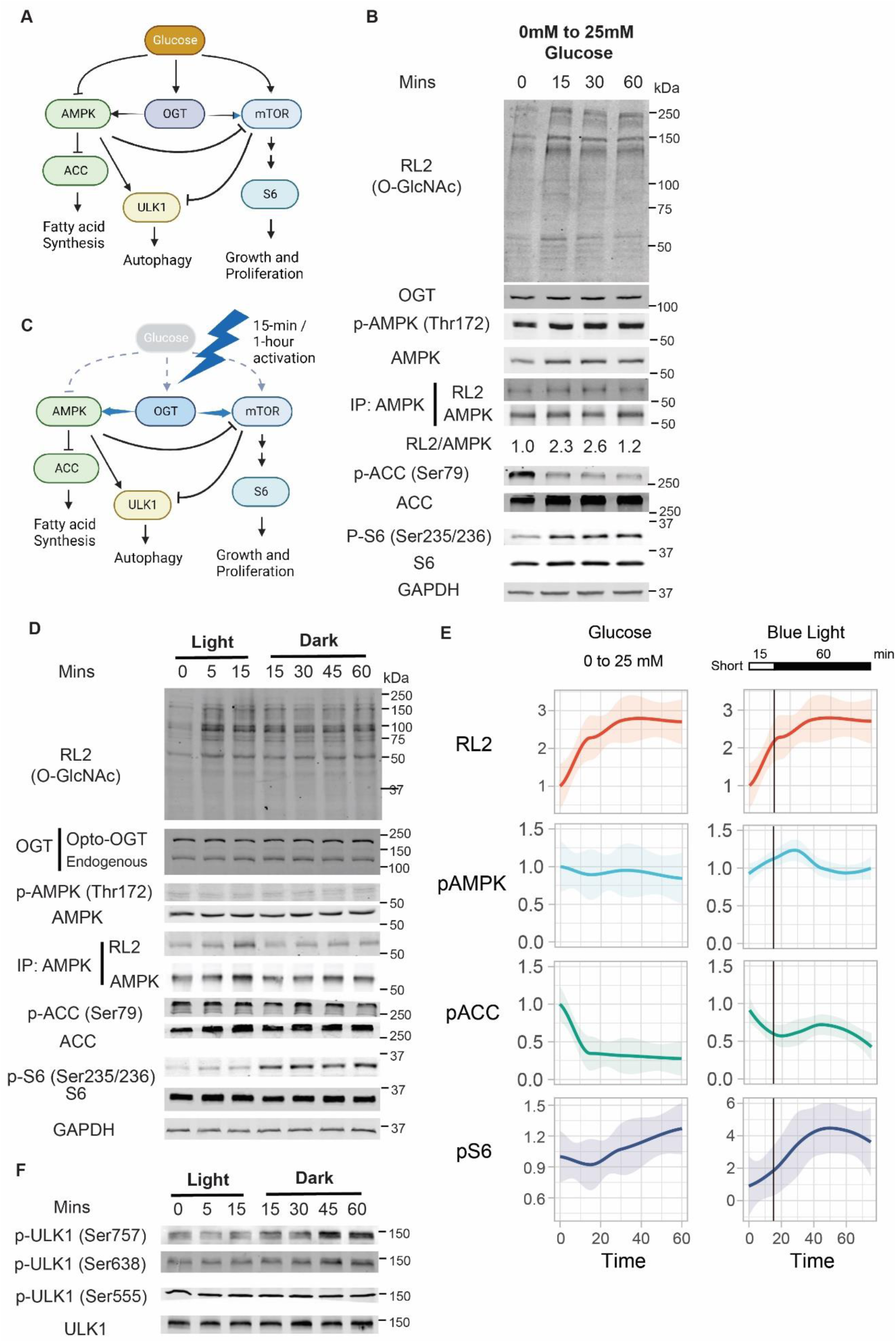
Optogenetic activation of OGT reveals temporal dependence for nutrient status sensing. (A) Schematic of key nutrient sensing pathways, OGT, AMPK and mTORC, working in concert to coordinate nutrient status to cellular responses. (B) Representative Western blots of key nutrient sensing pathways when HEK293T cells are subjected to glucose concentration changes from 0 mM to 25 mM. (C) Schematic of key nutrient sensing pathways, AMPK and mTORC, coordinates signaling input when Opto-OGT is activated for 15 minutes or an hour. (D) Representative Western blots of HEK293T cells transfected with OGT-mCherry-CRY2 undergoing 15 minutes of blue light. Key downstream signaling pathways are probed as labelled. (E) Quantification (n=2-3) of normalized phosphorylation signaling for Western blots depicted in (B) and (D). (F) Representative Western blots of phosphor-ULK1 when HEK293T cells transfected with OGT-mCherry-CRY2 underwent 15 minutes of blue light.

### Analysis of signaling crosstalk from light activation of Opto-OGT

It has been established that O-GlcNAcylation regulates diverse signaling pathways through crosstalk with phosphorylation^21^. Given that OGT has been reported to be a glucose sensor, and how O-GlcNAcylation changes are observed with glucose concentration changes, we asked if transient activation of OGT alone could result in similar signaling changes (**Figure 3C**). A 15-minute activation resulted in transient O-GlcNAcylation of AMPK, gradual phosphorylation of S6 and mTOR, while phosphorylation of AMPK remained unchanged, mimicking the effects of 0 mM to 25 mM glucose change (**Figures 3D, 3E, and S6B**).

We then observed the cellular responses arising from Opto-OGT activation with that observed from glucose concentration changes. These were reflected in autophagic signaling highlighted by ULK1 phosphorylation. In line with Opto-OGT-induced phosphorylation of mTOR, ULK1 phosphorylation at Ser757 by mTOR was increased, whereas there were no pronounced changes on two AMPK phosphorylation sites (Ser555 and Ser638) (**Figures 3F and S6A**).

### Analysis of AMPK, mTORC and O-GlcNAc signaling pathways in response to glucose level changes

We also performed glucose changes from 25 mM to 0 mM, and observed that O-GlcNAcylation of AMPK showed an prominent and persistent increase at 30 and 60 minutes (**Figure S6C**). In this regime, O-GlcNAc levels slightly decreased with increased phosphorylation of AMPK and ACC and decreased phosphorylation of S6 as expected (**Figure S6D**). For cells initially incubated at 2 mM glucose, similar trends are observed for ACC, AMPK and S6 phosphorylation though reduction of ACC phosphorylation became more delayed. Similar trends are also observed when the glucose concentration was changed from 25 mM to 2 mM (**Figure S6D**). When we activated Opto-OGT for an hour, we observed AMPK phosphorylation, continual AMPK O-GlcNAcylation and no changes in S6 phosphorylation (**Figures S6E**). To ensure that the differences did not arise from light-induced effects, we performed blue light exposure of both 15 mins and 1 hour on untransfected cells and found that key signaling pathways probed, namely O-GlcNAcylation, AMPK phosphorylation and S6 phosphorylation, did not change over the time course (**Figure S7**). Together, these results suggest that OGT is a crucial modulator balancing two opposing glucose sensing pathways (mTORC and AMPK signaling), and this effect can be clearly dissected by Opto-OGT tool.

### Targeting of Opto-OGT at mitochondria selectively increases mitochondrial O-GlcNAc levels

CRY2 can not only undergo homo-oligomerization, but can form heterodimers with the crytochrome-interacting basic-helix-loop-helix 1 protein (CIB1) upon blue light stimulation^45^. We used this feature to recruit OGT to specific subcellular locations by generating the fusion proteins of the N-terminus truncated version of CIB1 (aa 1-170), CIBN, with specific cellular location markers. We designed CIBN-GFP-miro that is localized at the outer membrane of the mitochondria, which is expected to sequester OGT-mCh-CRY2 from the cytosol towards the mitochondria and form oligomers in a light-controlled manner (**Figure 4A**)^46^. Indeed, in the presence of CIBN-GFP-miro, OGT-mCh-CRY2 translocated to the mitochondria upon blue light stimulation (**Movie S2, Figure S8A**). Colocalization analysis reveals the recruitment of OGT-mCh-CRY2 at CIBN-GFP-miro within 15 seconds of blue light illumination (**Figure S8B**).

**Figure 4:**
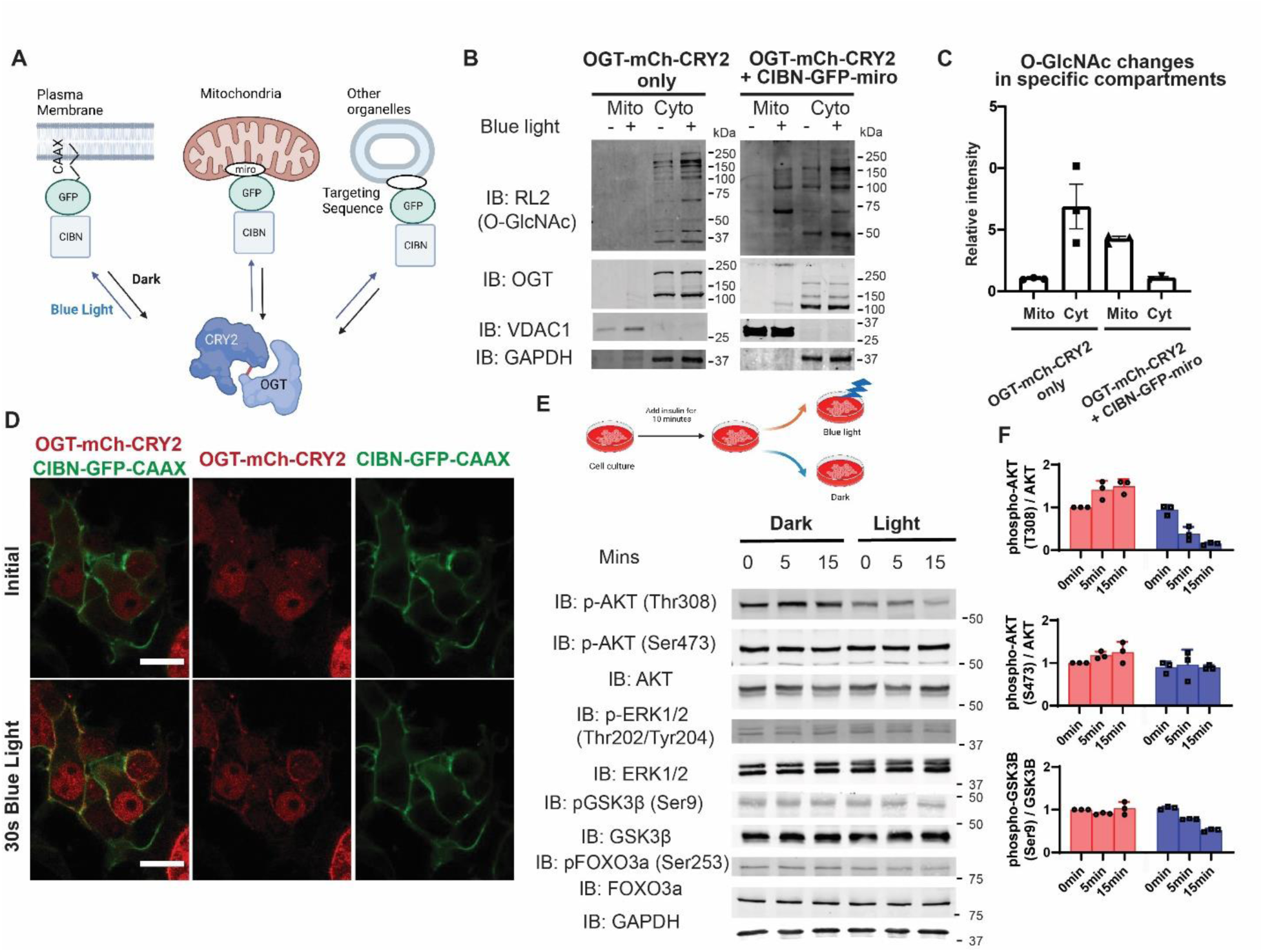
Optogenetic activation of OGT *via* CIBN-CRY2 recruitment results in subcellular compartment-specific increase of O-GlcNAcylation. (A) Schematic of light-controlled activation of Opto-OGT at different subcellular organelles and regions. This is achieved *via* CRY2-CIBN interaction with oligomerized OGT-mCherry-CRY2 localized to CIBN anchored to different organelles within the cell. (B) Western blot of subcellular fractionation experiment depicting RL2 (Global O-GlcNAc), OGT, VDAC1 and GAPDH of cells containing OGT-mCherry-CRY2 with or without CIBN-GFP-miro and subjected to dark or blue light. (C) Quantification (n=3) of normalized global O-GlcNAc levels relative to VDAC and GAPDH for Western blots depicted in (C). (D) Representative images of HEK293T cells transfected with OGT-mCherry-CRY2 and CIBN-GFP-CAAX from dark to blue light exposure. CIBN-GFP-CAAX could be found to colocalize with OGT-mCherry-CRY2 after 30 seconds of blue light exposure. Scale bar = 20 µm (E) Western blot of HEK293T cells transfected with OGT-mCherry-CRY2 and CIBN-GFP-CAAX undergoing 0, 5 and 15 minutes of blue light or dark treatment after insulin induction for 10 minutes. Key downstream signaling pathways are probed as labelled. (F) Quantification (n=3) of normalized phosphorylation signaling for Western blots depicted in (E).

We then tested whether the recruitment of Opto-OGT selectively increases mitochondrial O-GlcNAc levels. Subcellular fractionation experiments revealed that in the cells transfected with OGT-mCh-CRY2 alone, the O-GlcNAc levels were barely detectable in the mitochondria but had a 6-fold increase in the cytosol after 30 minutes of blue light illumination. In contrast, in the cells co-transfected with OGT-mCh-CRY2 and CIBN-GFP-miro, the O-GlcNAc levels showed a 4-fold increase in the mitochondria after blue light exposure but only a slight increase in the cytosol (**Figures 4B and 4C**). We also tested the effect of Thiamet-G, an OGA inhibitor, on O-GlcNAcylation in different subcellular compartments. Surprisingly, the O-GlcNAc levels in the mitochondria decreased after 30 minutes of Thiamet-G addition and returned to the basal level after 60 minutes; however, the O-GlcNAc levels in the cytosol increased steadily from 30 minutes to 120 minutes (**Figure S9**). Therefore, in contrast to a promiscuous effect of the OGA inhibitor, light-induced targeting of Opto-OGT at the mitochondria can modulate O-GlcNAcylation at the organelle level.

### Targeting of Opto-OGT at plasma membrane confirms the role of OGT in attenuating insulin signaling

The strategy used to target the mitochondria can be adapted to other cellular components. Our previous studies have shown that phosphoinositide-mediated recruitment of OGT to the plasma membrane selectively inhibits AKT phosphorylation at Thr308 and attenuates insulin signal transduction^47^. Here we utilized the prenylation tag, CAAX, to install the CIBN motif at the plasma membrane to enable Opto-OGT recruitment (**Figure 4A**). Upon 30 seconds of blue light stimulation, OGT-mCh-CRY2 translocated to the plasma membrane in HEK293T cells co-transfected with OGT-mCh-CRY2 and CIBN-GFP-CAAX (**Figure 4D, Movie S3**). Blue light exposure resulted in a rapid increase in global O-GlcNAcylation in these cells which persisted at 30 minutes (**Figure S10A**). We tracked the signaling pathways perturbed by light-induced recruitment of OGT towards the plasma membrane after 10 minutes of insulin introduction (**Figure 4E**). AKT phosphorylation at Thr308 was downregulated after 15 minutes of light exposure whilst Ser473 phosphorylation remained unchanged (**Figures 4E and 4F**). As a consequence, AKT-mediated phosphorylation of GSK3β at Ser9 and phosphorylation of FOXO3a at Ser253 were downregulated at 15 minutes. ERK1/2 phosphorylation was left unperturbed at 15 minutes (**Figures 4E and 4F**). To confirm if OGT recruitment to the plasma membrane indeed drives O-GlcNAcylation of AKT and thus attenuates its phosphorylation status, we performed immunoprecipitation of AKT and found significant O-GlcNAcylation on AKT upon blue light activation of OGT at the plasma membrane (**Figure S10B**). These results agree with previous reports that OGT translocation to the plasma membrane is key in attenuating insulin signal transduction, with the downregulation of AKT/GSK3β/FOXO signaling occurring within 15 minutes^47^.

## Discussion

In the realm of cell signaling, information is relayed from external cues to intracellular effectors in a coordinated manner. One critical external factor is the availability of nutrients, where intracellular sensors play a prominent role in activating cellular effectors. O-GlcNAcylation, mediated by the enzyme OGT, has been proposed as a nutrient sensor, but many questions remain unanswered. For example, how does OGT interact with other well-known nutrient sensors such as AMPK and mTORC ^43,48^? Moreover, OGT is known to be activated in both nutrient-rich and nutrient-deprived conditions, but the mechanisms underlying these dual signaling functions remain unclear ^3^. To gain insights into the role of O-GlcNAc in nutrient sensing, it is crucial to systematically modulate the dynamics of OGT activity and examine the resulting outputs in a rigorous manner^21^.

Efforts have been undertaken to modulate OGT signaling activities in order to gain insights into the role of O-GlcNAcylation in cellular regulation. Nutrients and hormones have been applied to increase O-GlcNAcylation and such approaches are key to elucidate its involvement in regulating insulin action. Genetic approaches, such as overexpression and silencing, as well as small molecule inhibitors, have been developed to perturb O-GlcNAcylation and impact cellular development. However, these approaches have limitations. Nutrients and growth factors, due to their tight binding, are not easily displaced and activate multiple signaling pathways simultaneously, making it challenging to isolate the role of O-GlcNAcylation. Genetic approaches are confounded by compensatory mechanisms that regulate O-GlcNAc levels ^49,50^. Furthermore, currently available inhibitors for OGT and OGA lack specificity and potency at the subcellular level, limiting their ability to dissect organelle-specific roles of O-GlcNAcylation. Therefore, the precise and spatiotemporal control of OGT activation offered by Opto-OGT is crucial in understanding the functional consequences of dynamic O-GlcNAcylation.

In this paper, we have demonstrated that glucose concentration changes are registered by OGT, with concomitant changes in AMPK O-GlcNAcylation, ACC phosphorylation and S6 phosphorylation. However, due to the likelihood of multiple events occurring simultaneously and our inability to capture other potential signaling events, it is challenging to determine if OGT is indeed the primary glucose sensor. We confirmed this notion through our findings that a 15-minute bout of Opto-OGT activation results in similar signaling events as observed with changes in glucose concentration from 0 mM to 25 mM. Interestingly, an extended regimen of 1-hour OGT activation shows opposite signaling patterns. This may suggest that OGT not only acts as a glucose sensor, but also as a nutrient shortage and stress sensor by affecting AMPK O-GlcNAcylation and subsequent signaling activity. Previous studies have established that manipulation of O-GlcNAcylation affects energy usage and AMPK activity in both nutrient-rich and nutrient-poor conditions ^9,43,51,52^. Our results provide insights into the potential mechanisms by which AMPK O-GlcNAcylation may be driven by OGT activation at different temporal regimes, ultimately influencing its phosphorylation status and activity.

Using Opto-OGT, we are also able to gain the understanding of the impact of OGT translocation to different organelles on cellular signaling and physiology. Activation of O-GlcNAc signaling at the mitochondria and plasma membrane is driven by the recruitment of OGT-mCherry-CRY2 to CIBN-miro and CIBN-CAAX, respectively. It has been demonstrated that OGT could be translocated to plasma membrane to regulate insulin signaling^47^ However, this is the first direct demonstration of how OGT activation at the plasma membrane leads to lowering of phosphorylation levels of AKT Thr-308. Study of the role of OGT at mitochondria has also caught attention recently within the field. While mitochondrial OGT has been known to impact mitochondrial activity and function, nucleocytoplasmic OGT may also O-GlcNAcylate mitochondrial proteins such as TOM70^53^. Thus, the Opto-OGT tool could be directed to O-GlcNAcylate proteins located at the outer membrane of the mitochondria to study these related signaling events.

The methodology we have undertaken in designing Opto-OGT is based on the local aggregation of full-length OGT, which results in its activation with high spatiotemporal control. Previous reports have cited the importance of multimerization of OGT in influencing the O-GlcNAcylation level of proteins, where TPR domains are involved in driving the association of different OGT proteins and the binding affinity of UDP-GlcNAc may be affected^54–57^. In this work, we demonstrated that the TPR domain is required for Opto-OGT action, even though the oligomerization of OGT is tuned by cryptochromes. We hypothesize that the TPR domains are potentially important for substrate recognition and binding, on top of their endogenous roles of being involved in multimerization of OGT. Thus, the mechanism underlying the workings of Opto-OGT could be due to the photo-uncaging of OGT with blue light which leads to local concentration increase of functioning OGT. Given that O-GlcNAcylation is controlled by a pair of enzymes – the writer being OGT and the eraser being OGA, local concentrations of OGT and OGA become an important part of the chemical equilibrium^21^. A unique property regarding Opto-OGT is that its substrate binding pocket is concealed in the dark state. Blue light is likely to cause conformational changes to cryptochrome which leads to its oligomerization while exposing the substrate binding site of Opto-OGT, allowing it to be active only under blue light. This results in Opto-OGT possessing no detectable background activity at effective concentrations and its activity is reversible and tunable.

There have been similar activities focusing on using light to probe the role of post-translational modifications, mostly via photo-uncaging^58^. Specifically, a recent attempt to construct a light-activated OGT toolkit was centered on replacing the catalytically essential lysine to a photocaged one. This resulted in an inactivated form of OGT which could be activated rapidly by photo-uncaging^59^. While the concept may be similar, our system possesses a few additional advantages. Firstly, the Opto-OGT is reversible and could be turned on/off at ease. Secondly, we utilize minimal intensity of blue light that does not cause phototoxicity or the increase of other signaling pathways in cells, thus allowing for dissection of signaling pathways. Thirdly, our system allows organelle level of control in inducing O-GlcNAcylation.

Recently, nanobody-based and aptamer-based approaches have been used to control the O-GlcNAc status of specific proteins^60–62^. We envision that the Opto-OGT tool could be similarly adapted to control protein-specific O-GlcNAcylation. Moreover, an orthogonal approach, for instance optogenetic methods with red light^63^, can be utilized to control OGA activity. The combinatorial optogenetic control of both OGT and OGA can be applied as a powerful tool in dissecting the role of O-GlcNAc in cellular signaling and function.

## Online Methods

### Cell culture and transfection

COS-7 and HEK293T cells were cultured in DMEM medium (Gibco) supplemented with 10% fetal bovine serum (Gibco) and 1% Penicillin-Streptomycin (Gibco). Lipofectamine 2000 (Thermo Fisher Scientific) was used to transfect the cells with the stated DNA plasmids according to the manufacturer protocol. Transfected cells were allowed to express the relevant proteins for 36 hours in complete culture medium under dark conditions before light illumination.

### Plasmid construction

The CIBN-GFP-CAAX, CRY2high-mCherry and CRY2low-mCherry plasmids were purchased from Addgene (Plasmid #26867, #104063 and #104065, respectively). In this work, we refer CIBN as a truncated domain of CIB1 containing amino acids 1 to 170. CRY2 refers to a modified version of CRY2 containing amino acids 1 to 498 with E490R modification. CIBN-GFP-Miro was constructed as previously described^46^.

For the plasmids OGT-mCherry-CRY2, OGT-mCherry-CRY2low, OGT(K852M)-mCherry-CRY2, sOGT-mCherry-CRY2 and CRY2-mCherry-Flag-OGT, the list of primers and methods used are recorded in **Table S1**.

### Light stimulation

For light illumination experiments, a blue LED array was constructed as adapted from an earlier report^30^. The array comprises of 24 blue LEDs (LED450L, Thorlabs). The light intensity at the height (12 cm) where the cell culture plate is placed was measured by a power meter (Newark, 1931-C) to be at 200 µW/cm^2^ with an absolute measurement uncertainty of less than 5%. The working distance of 12-cm is to ensure uniform blue light intensity across the surface of the cell culture dishes. The design of the homemade LED device is described in **Figure S11**.

### Western blot

For Western blot, the HEK293T cells were exposed to either light, complete darkness or light with dark recovery as described. After the relevant treatment, the cells were immediately washed in phosphate saline buffer before being subjected to RIPA buffer lysis supplemented with 20 µM Thiamet-G (Sigma Aldrich), cOmplete protease inhibitor cocktail tablet (Roche) and PhoSTOP (Roche). The cells were then agitated in 4°C for 30 minutes and centrifuged at 15000rpm for 20 minutes. The supernatant was then collected with protein concentration quantitated with BCA assay (ThermoFisher) on a plate reader (Tecan).

Subsequently, Lamelli buffer was added into the samples and subjected to SDS-PAGE. The gels were then transferred to a PVDF membrane using an eBlot system (Genscript) under the “SHORT” setting. The PVDF membrane is then blocked with 5% BSA in TBS containing 0.1% Tween-20 for an hour before addition of the relevant primary antibodies in blocking buffer for overnight. The blots are then washed in TBS containing 0.1% TBST for three times before being incubated in the relevant secondary antibodies and imaged on an Odyssey CLx (LI-COR).

The list of antibodies can be found in **Table S2**.

### Proteomics sample preparation

Samples for proteomics analysis were prepared as such: Anti-FLAG M2 affinity gel A2220(Sigma Aldrich) beads were twice washed with TBS, vortexed and centrifuged at 8000g for 1 minute. The clarified lysates, at about 500ug, were then added to the washed beads with a combined volume of 1ml. The samples were incubated overnight on a rotator at 4°C for binding. Subsequently, the samples were washed with phosphate-buffered saline (PBS) twice and supernatant was removed completely. Beads were then resuspended in 50% (v/v) trifluoroethanol (TFE) in 50 mM triethylammonium bicarbonate (TEAB), pH 8.5 containing 10 mM final concentration of tris(2-carboxyethyl)phosphine (TCEP) and incubated for 20 min at 55 °C for disulfide bridge reduction. Samples were cooled to 25 °C and alkylated with 55 mM 2-chloroacetamide (CAA) in the dark for 30 min, followed by on-bead digestion with endoproteinase LysC (2.5 µg final amount) for 3 h and subsequently by trypsin (2.5 µg final amount) at 37 °C overnight. Once completed, beads were removed and the peptides were transferred to new tubes. Digestion was terminated by adding 1% (v/v) final concentration of trifluoroacetic acid (TFA) to the samples, followed by desalting using C18 StageTips. Desalted peptides were dried by centrifugal evaporation, resuspended in 25 µl of 100 mM TEAB, pH 8.5, and individually labelled using isobaric tandem mass tags (TMT10-plex, Thermo Fisher Scientific) at 25 °C overnight. After labelling was completed, the reaction was quenched by addition of 10 µl of 1 M Tris, pH 8 into each tube before pooling the samples into a new low-binding 1.5-ml microfuge tube. Pooled sample was desalted and fractionated on a self-packed spin column containing C18 beads (Dr Maisch GmbH) using 14%, 18%, 21%, 24%, 27%, 32%, and 60% acetonitrile in 10 mM ammonium formate, pH 10 as the step gradients. Fractions were dried by centrifugal evaporation and further washed and dried twice by addition of 60% acetonitrile in 0.1% formic acid to further remove residual ammonium formate salts.

### Tandem mass spectrometry analysis

Dried fractions were resuspended in 10 µl of 2% (v/v) acetonitrile containing 0.06% (v/v) trifluoroacetic acid and 0.5% (v/v) acetic acid and transferred to an autosampler plate. Online chromatography was performed in an EASY-nLC 1000 (Thermo Fisher Scientific) liquid chromatography system using a single-column setup and 0.1% formic acid in water and 0.1% formic acid in 99% acetonitrile as mobile phases. Fractions were injected and separated on a reversed-phase C18 analytical column (Easy-Spray, 75 µm inner diameter × 50 cm length, 2 µm particle size, Thermo Fisher Scientific) maintained at 50 °C and using a 2-33% (v/v) acetonitrile gradient over 55 min, followed by an increase to 45% over the next 5 min, and to 95% over 5 min. The final mixture was maintained on the column for 4 min to elute all remaining peptides. Total run duration for each sample was 70 min at a constant flow rate of 300 nl/min.

Data were acquired using an Q Exactive HFX mass spectrometer (Thermo Fisher Scientific) using data-dependent mode. Samples were ionized using 2.1 kV and 300 °C at the nanospray source. Positively-charged precursor signals (MS1) were detected using an Orbitrap analyzer set to 60,000 resolution, automatic gain control (AGC) target of 3,000,000 ions, and maximum injection time (IT) of 50 ms. Precursors with charges 2-7 and having the highest ion counts in each MS1 scan were further fragmented using higher-energy collision dissociation (HCD) at 37% normalized collision energy. Fragment signals (MS2) were analysed by the Orbitrap analyzer at a resolution of 45,000, AGC of 100,000 and maximum IT of 100 ms. Precursors used for MS2 scans were excluded for 30 s to avoid re-sampling of high abundance peptides. The MS1–MS2 cycles were repeated every 20 MS2 scans until completion of the run.

### Proteomics data analysis

Proteins were identified using Proteome Discoverer™ (v2.4, Thermo Fisher Scientific). Raw mass spectra were searched against human primary protein sequences retrieved from Swiss-Prot (11 June 2019). Carbamidomethylation on Cys and TMT6-plex on Lys and N-terminus were set as a fixed modification; deamidation of asparagine and glutamine, acetylation on protein N-termini, and methionine oxidation were set as dynamic modifications for the search. Trypsin/P was set as the digestion enzyme and was allowed up to three missed cleavage sites. Precursors and fragments were accepted if they had a mass error within 10 ppm and 0.06 Da, respectively. Peptides were matched to spectra at a false discovery rate (FDR) of 1% (strict) and 5% (relaxed) against the decoy database and quantitated using TMT6-plex method. Search result was exported and further processed for comparative analysis. The results were normalized and checked if the counts were significantly different in both sets of data.

### Subcellular Fractionation

For subcellular fractionation, cells are lysed and resuspended using subcellular fractionation buffer (20mM HEPES (pH7.4), 10mM KCl, 2mM MgCl_2_, 1mM EDTA, 1mM EGTA, 1mM DTT, Phosphatase Inhibitor Mini Tablets (Pierce) and Complete Protease Inhibitor Cocktail (Roche)) before being incubated on ice for 15 minutes. Cells are further lysed by passing the suspension through a 27-gauge needle and left on ice for 30 minutes. Samples are then centrifuged at 720g (3000rpm) for 5 minutes (at 4°C) to separate the nuclear pellet from supernatant containing the mitochondria and cytosol. Supernatant containing mitochondria and cytosol is centrifuged at 10,000g (8000rpm) for 5 minutes (at 4°C), separating the mitochondrial pellet from the supernatant containing the cytosol. Supernatant containing the cytosol is sonicated briefly and stored at -80°C for subsequent protein analysis using Western blot. To wash the mitochondrial pellet, 500ul of subcellular fractionation buffer is added and the pellet is dispersed using a 25-guage needle; the suspension is then spun down at 3000 rpm for 10 minutes (at 4°C) and pellet containing nuclei is discarded while the supernatant containing mitochondria is kept and spun down at 8000rpm for 10 minutes (at 4°C). Supernatant is discarded and mitochondrial pellet is resuspended in NP-40 lysis buffer with 0.1% SDS. The mitochondria are lysed by sonicating the suspension briefly whilst on ice. Final mitochondrial fraction is stored at -80°C for subsequent protein analysis using Western blot.

### Fluorescent microscopy on fixed cells

COS-7 cells transfected with OGT-mCherry-CRY2 and subjected to either light or dark treatment were first washed with PBS and then fixed with 4% paraformaldehyde for 20 minutes at room temperature. A second round of gentle washing ensues before the cells are subjected to 0.5% of Triton X-100 in PBS for 15 minutes at room temperature. 2% bovine serum albumin in PBS is then added and incubated for an hour at room temperature. The cells were then incubated with primary antibodies (1:500 for OGT and 1:100 for RL2), washed three times with PBS before the corresponding secondary antibodies addition at 1:1000 dilution. The cells were then stained with DAPI before being imaged with LSM 880 Airyscan Confocal Microscope (Zeiss). The list of antibodies used can be found in **Table S2**.

### Live cell imaging

For live cell imaging noting on the reversible oligomerization of OGT-mCherry-CRY2, transfected cells were recorded at every 5-second interval using LSM 880 Airyscan Confocal Microscope equipped with a temperature and carbon dioxide-controlled environmental chamber for extended live cell imaging. Each pulse of blue light is supplied as a 100-ms pulse at 488nm.

### Quantiative puncta measurement

The raw images are first split into single channel images, with the red channel only taken into account. Using imageJ, background correction is applied and set for a uniform threshold before the images are analyzed by the Speckle Counting pipeline available on CellProfiler^64^. In this pipeline, we first identified cells with the “IdentifyPrimaryObjects” module by using the maximal intensity image as the input image. Within the module, “Global Otsu Thresholding” was selected to perform two-class thresholding. Subsequently, “EnhanceOrSuppressFeatures” module was employed to select for the particles and “MaskImage” was then used to generate a mask to restrict particle counting within each cell. To identify the individual particles located within each image, the image previously derived from “MaskImage” module was measured for each of their individual diameter. Lastly, “RelateObjects” was used to directly relate counts to individual cells.

### Colocalization analysis

The raw images are first split into single channel images, with the red and green channels taken into account. Using imageJ, background correction is applied and set for a uniform threshold before the images are analyzed by the Measure Colocalization pipeline available on CellProfiler^64^. 15% threshold is set as the threshold for maximum intensity for the images.

### Simulation of protein structures

Simulation of fusion protein structure and heterodimerization of CRY2 and OGT was done using Alphafold2^39^, utilizing Colabfold method previously described on the Google Colaboratory server^65^. Subsequent visual models were created using PyMOL 2.5 (Shroedinger). ClusPro analysis^66,67^ is also run based on protein PDB structures 7D0N (for CRY2) and 7NTF (for OGT).

### Quantification and Statistical Analyses

The number of replicates, error bar representation and statistical significance are indicated in the respective figure legends. All statistical analyses were performed with the GraphPad Prism Software 8.4.3 (GraphPad Software Inc., La Jolla, CA, USA).

### Reporting summary

Further information on research design is available in the Nature Portfolio Reporting Summary linked to this article.

## Data availability statement

The mass spectrometry proteomics data is deposited to the JPOST repository with the dataset identifier JPST002053. Further information on the analyzed mass spectrometry data and AlphaFold simulation is available in the Supplementary Information. Source data is provided with this paper. The data generated and/or analysis during the current study are available from the corresponding author on reasonable request.

## Acknowledgements

This work was supported by the National Institutes of Health (R01DK089098, R01DK102648) and American Diabetes Association (1-19-IBS-119) to X.Y., Intramural support from the Agency for Science, Technology and Research (A*STAR) Biomedical Research Council core fund, A*STAR Strategic Research Program (the Brain-Body Initiative, iGrants call ID #21718), A*STAR Use-Inspired Basic Research Award, and National Research Foundation Competitive Research Program (CRP) (NRF-CRP23- 2019-0004) to W.H. Q.O. gratefully acknowledges support from the A*STAR YIRG grant (OFYIRG19nov-0045). We thank Dr. Pei Ling Chia and Ms Haitong Mao for proof-reading of the manuscript.

## Contributions

Q.O., W.H. and X.Y. conceived the project. Q.O., Y.L. and X.Y. designed the experiments. Q.O., R.L., C.G., S.E.C., V.K., J.Y., T.T., K.S., S.L.L., L.C.W and S.G.L. performed the experiments. Q.O., C.G., Y.L., V.K., T.T., L.C.W and S.G.L. analyzed the data. R.L., S.E.C., R.D. and R.S. offered technical advice. Q.O., A.B., W.H. and X.Y. wrote the paper with input from all authors.

