## Supplementary Figures for "Spatiotemporal control of subcellular O-GlcNAc signaling using Opto-OGT"

**Supplementary Tables and Figures**

**Table S1**: List of primers used in this study

| Plasmid | Method | Primers Used: |
|---|---|---|
| OGT-mCherry-CRY2 | InFusion cloning with PCR-amplified CRY2high from CRY2high-mCherry plasmid cloned into linearized phOGT-Kozak-mcherry-2 X 3GSS | Forward: TCTTCAGGATCATCCatgaagatggacaaaaagactatagtttggt<br><br>Reverse:<br>attgcattcattttatgctgctccgatcatgatctg |
| OGT-mCherry-CRY2low | InFusion cloning with PCR-amplified CRY2low from CRY2low-mCherry plasmid cloned into linearized phOGT-Kozak-mcherry-2 X 3GSS | Forward:<br>TCTTCAGGATCATCCatgaagatggacaaaaagactatagtttggt<br><br>Reverse:<br>attgcattcattttagtcttcctcggttcttgaaatagct |
| OGT(K852M)-mCherry-CRY2 | InFusion cloning with OGT-mCherry-CRY2 as template | Forward:<br>TTGTATATGATTGACCCTTCTACTTTGCA<br><br>Reverse:<br>GTCAATCATATACAACTGATTAAAGTTACAGT |
| sOGT-mCherry-CRY2 | InFusion cloning with OGT-mCherry-CRY2 as template | Forward:<br>ACGCCACCATGCATTATAAGGAGGCTATTCGA<br><br>Reverse:<br>AATGCATGGTGGCGTCGACAGATCTG |

| | | |
|---|---|---|
| CRY2-mCherry-OGT | Gibson cloning with PCR-amplified OGT from OGT-mCherry plasmid as insert cloned into linearized CRY2high-mCherry from CRY2high-mCherry-Raf1 | Forward (insert):<br>ccggactcagatctcgagtgATGGCGTCTTCCGTGGGC<br>Reverse (insert):<br>tagactcgagcggccgcttaTGCTGACTCAGTGACTTCAACAG<br>Forward (vector): TAAGCGGCCGCTCGAGTC<br>Reverse (vector): CACTCGAGATCTGAGTCCGG |

**Table S2**: List of antibodies used in this study

| Primary Antibody | Source | Identifier / Added ratio in western blots |
|---|---|---|
| OGT | Cell Signaling Technology | Cat #24083, 1:1000 |
| MGEA5 / OGA | Abcam | ab 124807, 1:1000 |
| RL2 | Abcam | ab 2739, 1:500 |
| GAPDH | Santa Cruz | sc-32233, 1:2000 |
| VDAC1 | Abcam | ab 154856, 1:1000 |
| AMPK | Santa Cruz | ac-25792, 1:1000 |
| Phospho-AMPK (Thr172) | Santa Cruz | sc-33524, 1:200 |
| ACC | Cell Signaling Technology | Cat: #3673S, 1:1000 |
| Phospho-ACC (Ser79) | Cell Signaling Technology | Cat: #3661S, 1:1000 |
| AKT (pan) | Cell Signaling Technology | Cat: #4691, 1:1000 |
| Phosphor-AKT (Ser473) | Cell Signaling Technology | Cat: #4051S, 1:1000 |
| Phosphor-AKT (Thr308) | Cell Signaling Technology | Cat #13038, 1:1000 |
| P44/42 MAPK (Erk1/2) | Cell Signaling Technology | Cat: #9102, 1:1000 |
| Phospho-p44/42 MAPK (Erk1/2) (Thr202/Tyr204) | Cell Signaling Technology | Cat: #9101, 1:1000 |
| P38 MAPK (D13E1) XP(R) | Cell Signaling Technology | Cat: #8690, 1:1000 |
| Phospho-p38 MAPK (Thr180/Tyr182) Antibody | Cell Signaling Technology | Cat: #9211, 1:1000 |
| SAPK/JNK | Cell Signaling Technology | Cat: #9252, 1:1000 |

| | | |
|---|---|---|
| Phospho-SAPK/JNK (Thr183/Tyr185) Antibody | Cell Signaling Technology | Cat: #9251, 1:1000 |
| GSK-3β (D5C5Z) | Cell Signaling Technology | Cat: #12456, 1:1000 |
| Phospho-GSK-3β (Ser9) (D85E12) | Cell Signaling Technology | Cat #5558, 1:1000 |
| FOXO3a | Cell Signaling Technology | Cat #9467, 1:1000 |
| Phospho-FOXO3a (Ser253) | Cell Signaling Technology | Cat #9466, 1:1000 |
| Phospho-ULK1 (Ser555) | Cell Signaling Technology | Cat #5869S, 1:1000 |
| Phospho-ULK1 (Ser638) | Cell Signaling Technology | Cat #14205S, 1:1000 |
| Phospho-ULK1 (Ser757) | Cell Signaling Technology | Cat #14202S, 1:1000 |
| ULK1 (D8H5) | Cell Signaling Technology | Cat #8054, 1:1000 |
| Secondary Antibody | Source | Added ratio in western blots |
| IRDye® 800CW Goat anti-Rabbit IgG (H + L) | LICOR | 1:10000 |
| IRDye® 680RD Donkey anti-Mouse IgG (H + L) | LICOR | 1:10000 |

| Secondary Antibody | Source | Added ratio in immunofluorescence experiments |
|---|---|---|
| Alexa Fluor® 488 goat-anti-mouse | Thermofisher | 1:1000 |
| Alexa Fluor® 647 goat anti-rabbit | Thermofisher | 1:1000 |

#### Supplemental Figure Legends

**Figure S1: AlphaFold Predictions of Opto-OGT tool in the dark state, in relation to Figure 1. (A)** Sequence coverage for multiple sequence alignment of Opto-OGT. **(B)** Predicted Local Distance Difference Test (LDDT) per position for multiple sequence alignment of Opto-OGT. (**C**) Plots of inter-chain predicted alignment error (inter-PAE) provided by AlphaFold2.

**Figure S2: Predictions of association of OGT and CRY2 in the dark state via ClusPro and AlphaFold, in relation to Figure 1. (A)** ClusPro simulation of PDB structures 7D0N (for CRY2) and 7NTF (for OGT) in the dark state reveals that CRY2 blocks access to OGT substrate binding pocket. **(B)** AlphaFold simulation of OGT and CRY2 in the dark state reveals that CRY2 blocks access to OGT substrate binding pocket. **(C)** Sequence coverage for multiple sequence alignment of Opto-OGT. **(D)** Predicted Local Distance Difference Test (LDDT) per position for multiple sequence alignment of Opto-OGT. **(E)** Plots of inter-chain predicted alignment error (inter-PAE) provided by AlphaFold2.

**Figure S3: Calibration and optimization of Opto-OGT construction, in relation to Figure 1. (A)** Quantification (n=6) of OGT and OGA expression relative to GAPDH for Western blots depicted in Figure 1C. **(B)** Representative Western blot of HEK293T cells transfected with mCherry-CRY2 or untransfected ones subject to 0, 10 and 30 minutes of blue light treatment.

**Figure S4: Opto-OGT activation leads to rise in local *O*-GlcNAc levels, in relation to Figure 1.** Representative images of COS-7 cells transfected with OGT-mCherry-CRY2 and undergoing either dark or blue light treatment. The yellow arrows indicate the colocalization of OGT-mCherry-CRY2 oligomers and O-GlcNAcylated sites. Scale bar = 20 µm.

**Figure S5: Calibration of experimental conditions in relation to Figures 1 and 2. (A)** Representative Western blot of HEK293T cells transfected with mCherry-CRY2 or untransfected ones subject to 0, 10 and 30 minutes of blue light treatment. The phosphorylation status of p38 and ERK1/2, which belong to the stress-sensitive MAPK pathways, are examined relative to GAPDH. **(B)** Representative Western blot of HEK293T cells grown on 10-cm dishes and transfected with different amounts of OGT-mCherry. The cellular O-GlcNAc levels are examined with respect to GAPDH.

**Figure S6: Representative Western blots depicting key glucose sensing pathway changes of HEK293T cells exposed to glucose concentration changes, in relation to Figure 3. (A)** Representative Western blots of MTOR, P70S6K and ULK pathways when HEK293T cells are subjected to glucose concentration changes from either 0 mM to 25 mM or vice versa. **(B)** Representative Western blots of HEK293T cells transfected with OGT-mCherry-CRY2 undergoing 15 minutes of blue light. mTOR and P70S6K pathways are probed as labelled. **(C)** Representative Western blots of O-GlcNAcylation of AMPK when HEK293T cells are subjected to glucose concentration changes from 25mM to 0mM. **(D)** Representative Western blots of key nutrient signaling pathways when HEK293T cells are subjected to glucose concentration changes from either 2 mM to 25 mM, 25mM to 2mM or 25mM to 0mM. **(E)** Representative Western blots of key nutrient signaling pathways when HEK293T cells transfected with Opto-OGT are subjected to 1 hour of blue light.

**Figure S7: The usage of blue light at 15 min or 1 hour does not affect O-GlcNAc, AMPK and S6 signaling on non-transfected cells, in relation to Figure 3.**

**Figure S8: Optical recruitment of activated OGT to mitochondria takes place within seconds in COS-7 cells. (A)** Representative images of COS-7 cells transfected with OGT-mCherry-CRY2 and CIBN-GFP-miro undergoing 0, 15, 60 and 300 seconds of blue light. **(B)** Quantification (n=5) of correlation coefficient of Opto-OGT and CIBN-GFP-miro.

**Figure S9: Usage of OGA inhibitor, Thiamet-G, does not allow subcellular control of O-GlcNAcylation, in relation to Figure 4.** Representative western blot of HEK293T cells subject to 0, 30, 60 and 120 minutes of 10μM Thiamet-G treatment (OGA inhibitor). The global O-GlcNAc levels (RL2) of mitochondria and cytosol are noted.

**Figure S10: Optical recruitment of activated OGT to plasma membrane results in increase in O-GlcNAc levels and O-GlcNAcylation of AKT. (A)** Western blot of HEK293T cells transfected with OGT-mCherry-CRY2 and CIBN-GFP-CAAX undergoing 0, 10 and 30 minutes of blue light. O-GlcNAc levels and GAPDH are probed. **(B)** Western blot of HEK293T cells under different conditions, with AKT immunoprecipitated and looking at O-GlcNAc levels of AKT.

**Figure S11: Diagram of homemade LED device.**

**Supplementary Movies**

**Movie S1:** Movie showing HEK293T cells transfected with OGT-mCherry-CRY2. The first 300 seconds are without blue light while in the next 300 seconds, the cells are supplied with blue light at a 100-millisecond pulse every 5 seconds. No blue light is illuminated from 600 seconds to 1800 seconds and blue light is reapplied from 1800 to 2400 seconds at a 100-millisecond pulse every 5 seconds.

**Movie S2:** Movie showing COS7 cells transfected with both OGT-mCherry-CRY2 (on the right panel) and CIBN-GFP-miro (on the left panel). Green and blue lights are continually supplied to the system at a 100-millisecond pulse every 15 seconds.

**Movie S3:** Movie showing HEK293T cells transfected with both OGT-mCherry-CRY2 (on the right panel) and CIBN-GFP-CAAX (on the left panel). Green and blue lights are continually supplied to the system at a 100-millisecond pulse every 5 seconds.

#### Related to Figure 1 – Figure S1

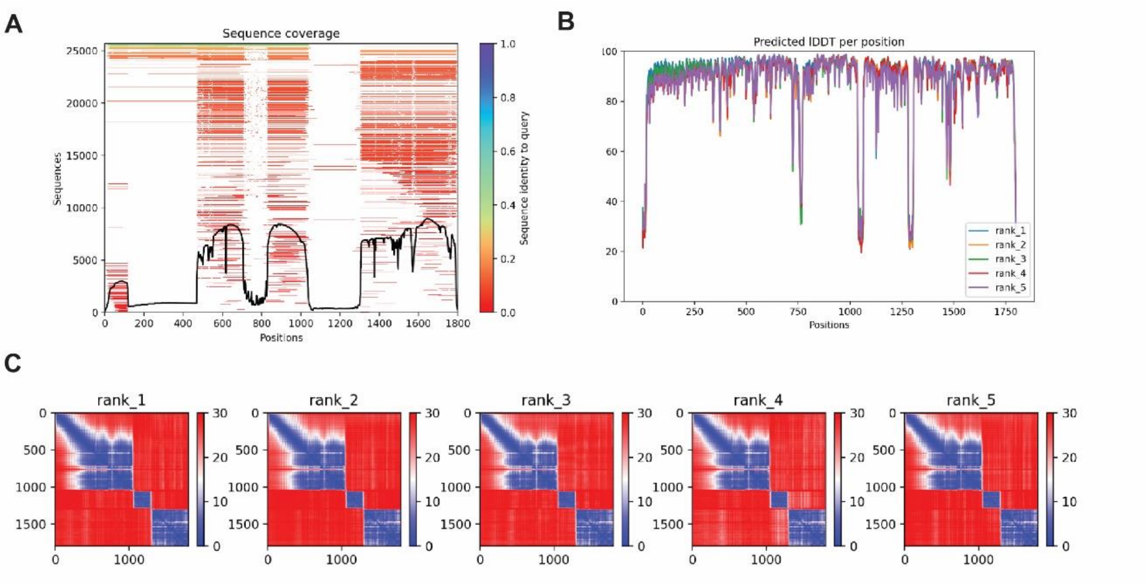

#### Related to Figure 1 – Figure S2

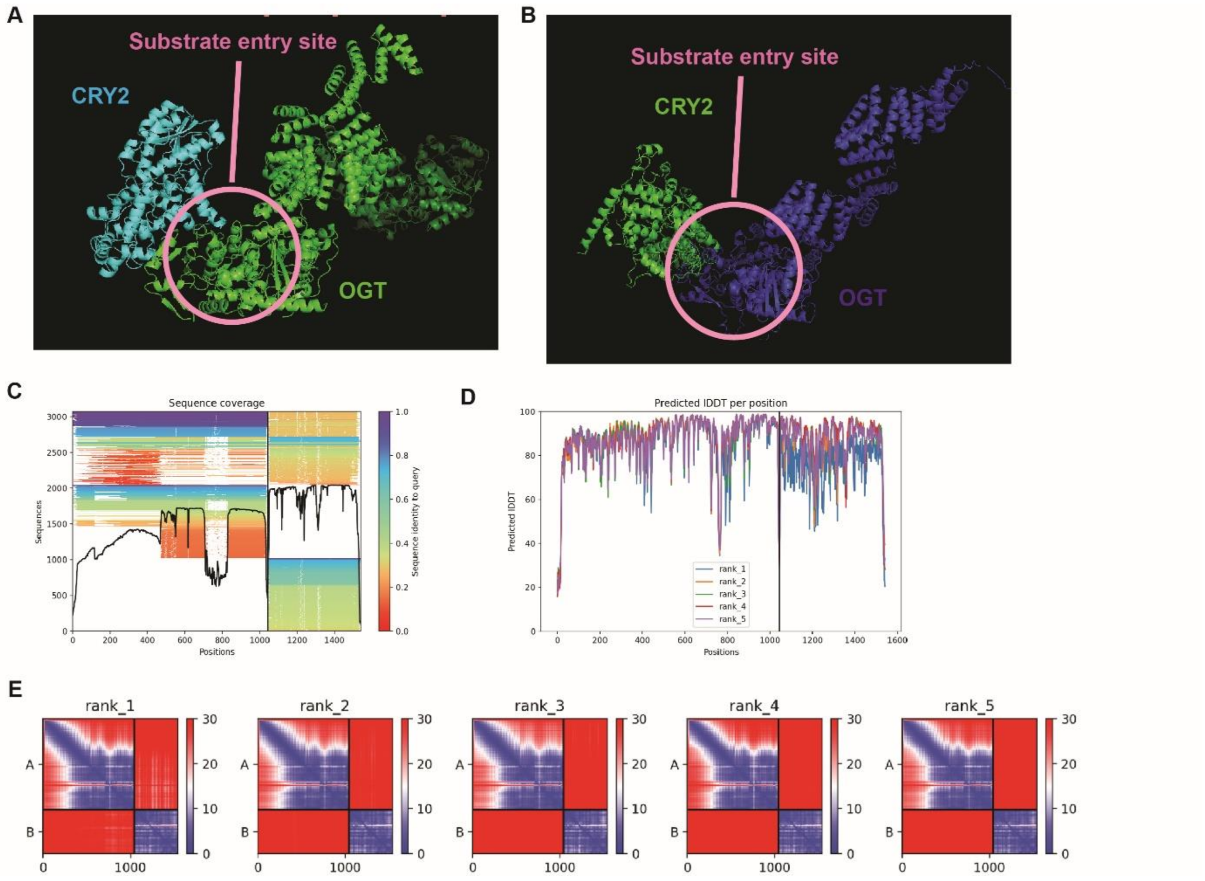

#### Related to Figure 1 – Figure S3

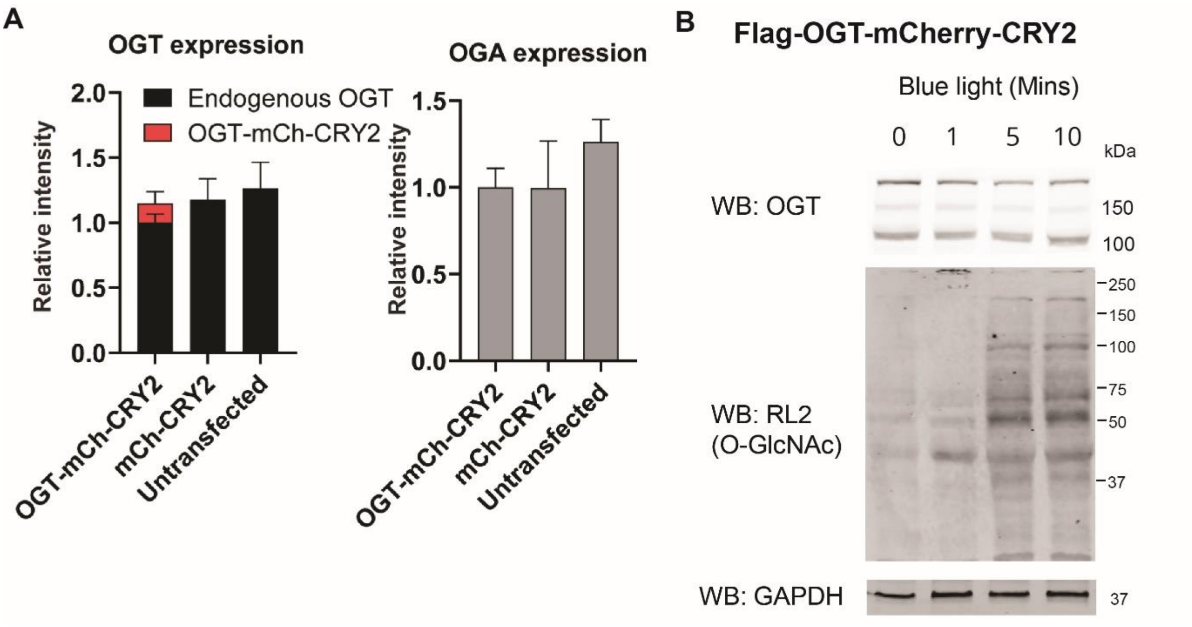

#### Related to Figure 2 – Figure S4

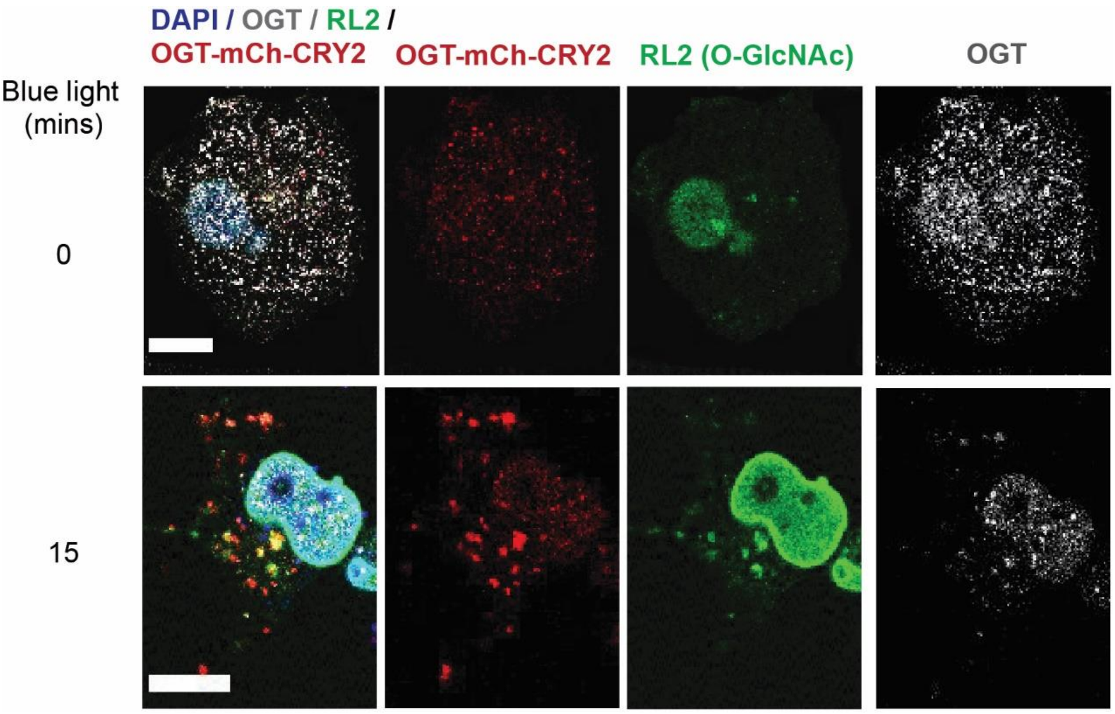

**Related to Figure 2 – Figure S5**

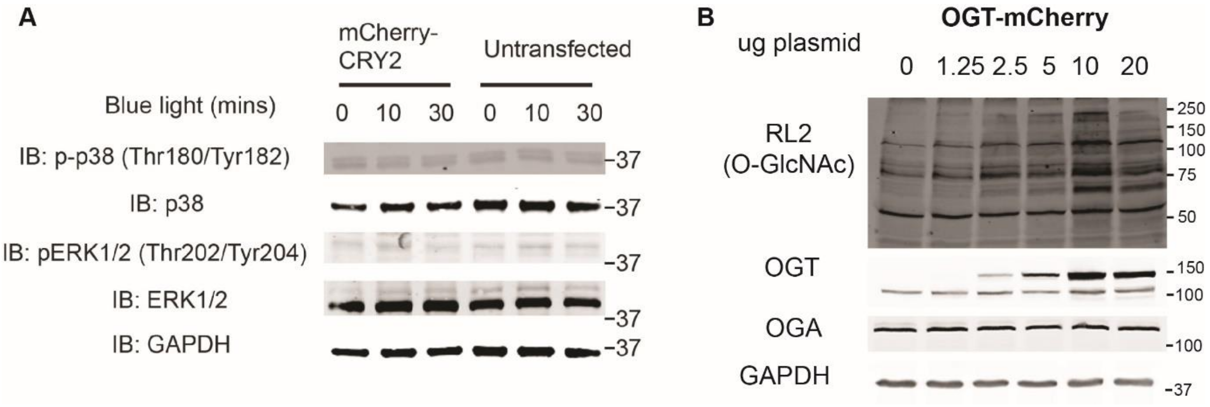

### Related to Figure 3 – Figure S6

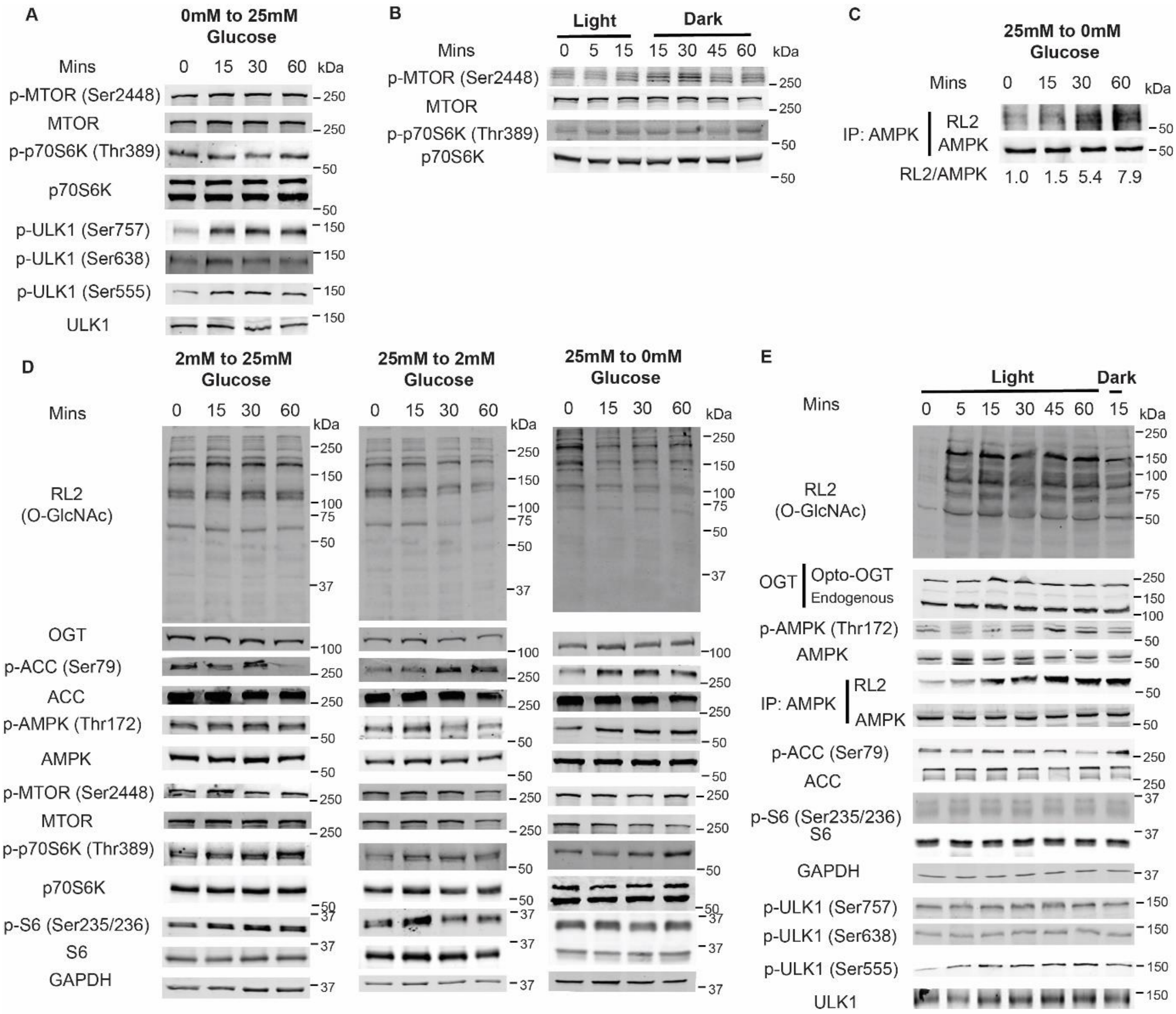

**Related to Figure 3 – Figure S7**

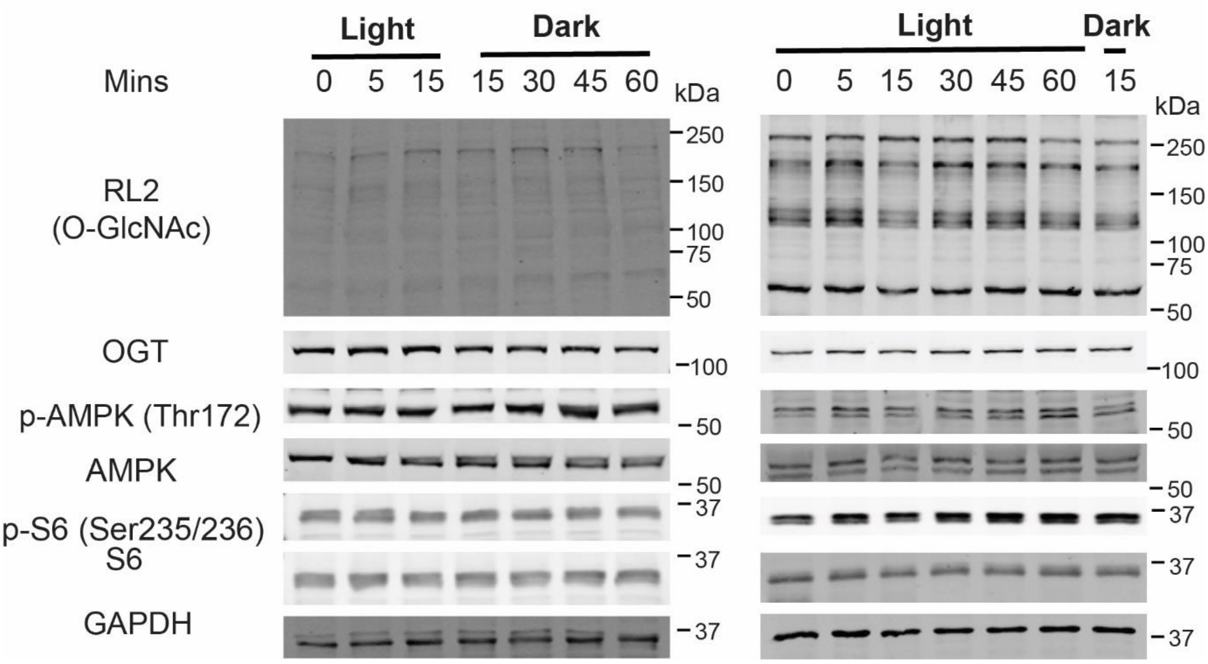

**Related to Figure 4 – Figure S8**

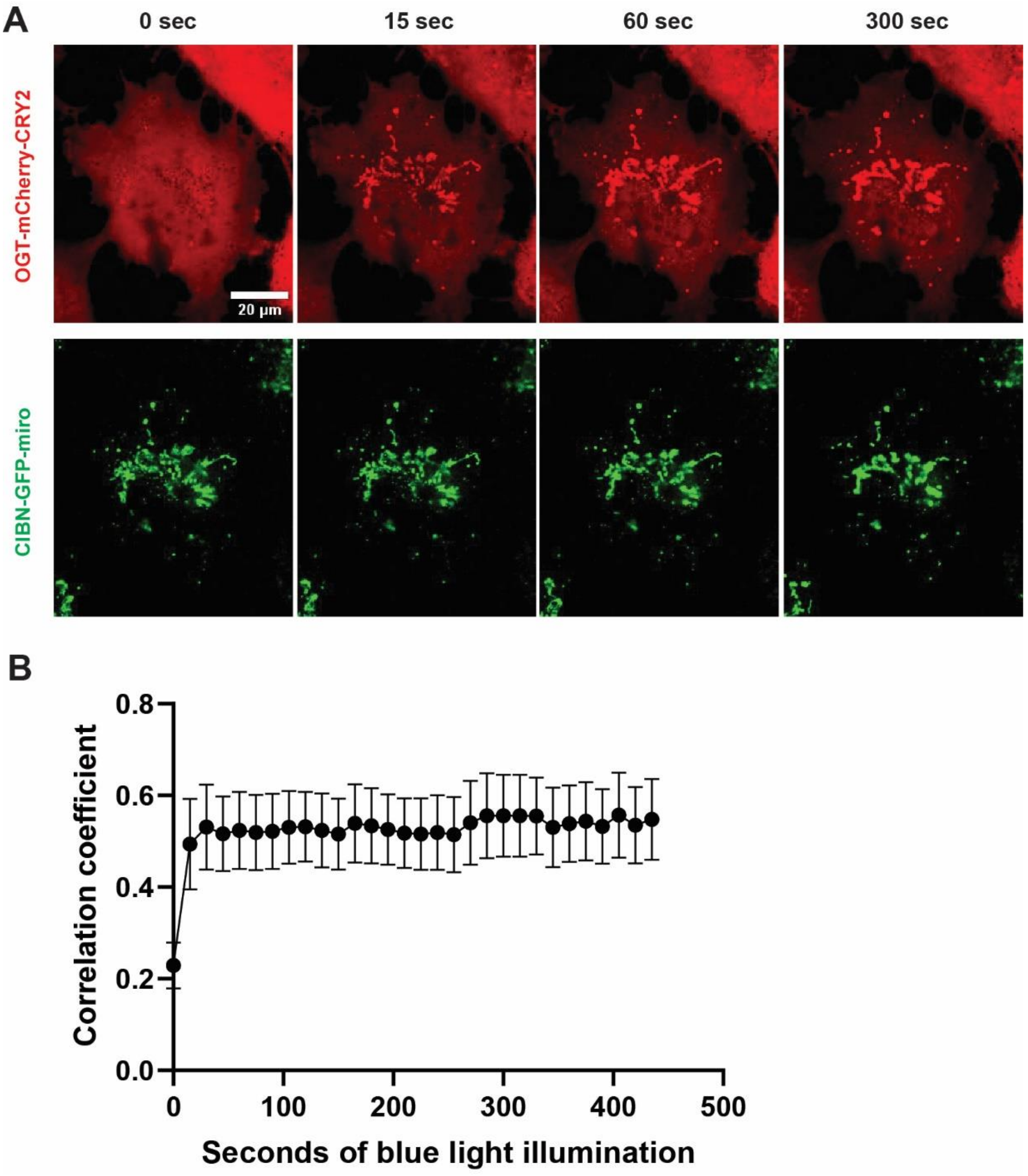

**Related to Figure 4 – Figure S9**

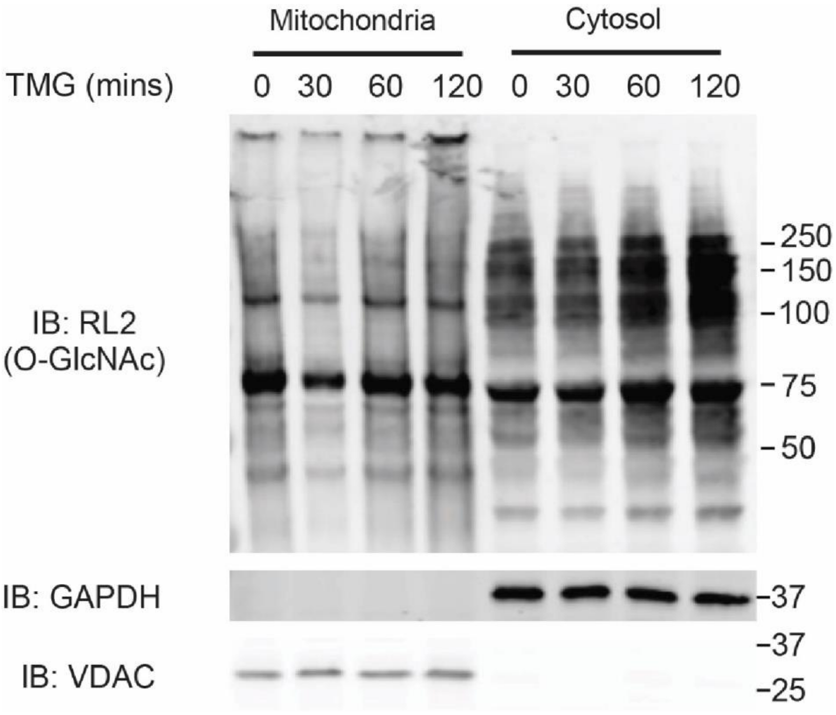

**Related to Figure 4 – Figure S10**

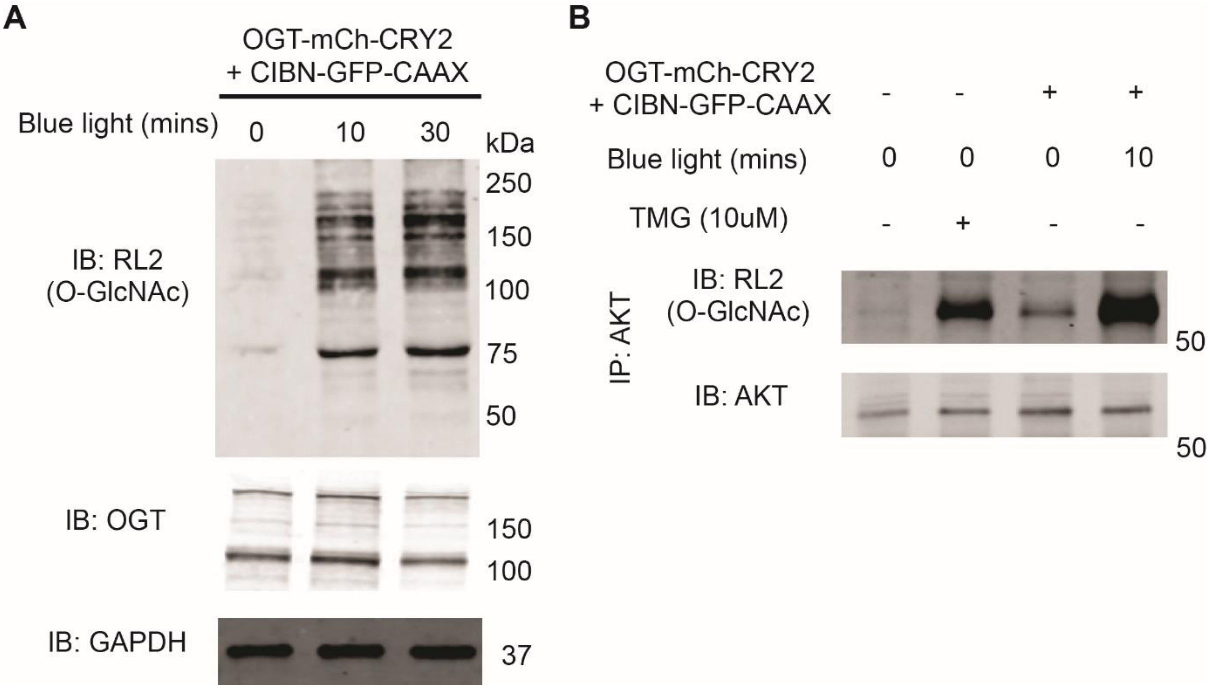

**Figure S11**

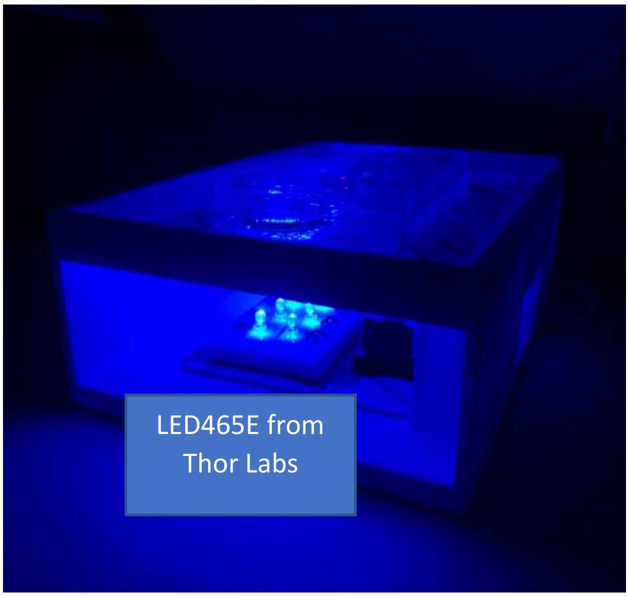
